# IRIS, a tool for the in-silico evaluation of mosquito control trial designs based on inundative releases

**DOI:** 10.1101/2025.11.05.686816

**Authors:** Jagadeesh Chitturi, Paulo C. Ventura, Allisandra G. Kummer, Chalmers Vasquez, Ethan SeRine, Megan D. Hill, Mattia Manica, Piero Poletti, John C. Beier, Keisuke Ejima, Michael Johansson, Stefano Merler, Hongjie Yu, John-Paul Mutebi, Maria Litvinova, André B.B. Wilke, Marco Ajelli

## Abstract

Mosquito control strategies based on the mass release of modified males, such as genetically modified mosquitoes (GMMs), aim to suppress wild populations by impairing reproduction. Evaluating these interventions requires resource-intensive field trials, but a lack of standardized implementation practices, particularly regarding release ratios of modified males to wild female mosquitoes and trial timing, has led to variable outcomes. This study’s objective is to propose a modeling tool for the “in-silico” simulation of trial designs before field implementation. To this aim, we developed an agent-based model of mosquito population dynamics. As a case study, we calibrated the model using 2019-2023 *Aedes aegypti* surveillance data from Miami-Dade County, Florida, and compared two GMM trial designs as illustrative examples. Our results show that depending on the implementation choices (e.g., trial start date and duration, release ratio), trials yield highly variable outcomes. For example, changing the start date while fixing all other implementation details can lead to effectiveness between 50% and 90%. Our findings suggest that “in-silico” simulation is a valuable tool for improving trial protocol design, allowing stakeholders to test strategies and reduce outcome uncertainty before committing to a fieldwork experiment.

## Introduction

Traditional insecticide-based strategies are the cornerstone of mosquito control [1–4]. However, they have limitations such as insecticide resistance [5–7] and potential environmental impacts and effects on biodiversity, ecosystems, and human health, regardless of the effectiveness on mosquito control [8]. Therefore, in past decades, a new generation of interventions has emerged, which are based on mass releases of irradiated sterile mosquitoes, *Wolbachia*-infected mosquitoes, or genetically modified mosquitoes (GMM) [9–18]. These strategies, here referred to as “inundative” mosquito control strategies, rely on mass releases of adult male mosquitoes that are either sterile or carrying a modified gene or endosymbiont, which impair reproduction [12,19]. However, implementing inundative strategies still requires rigorous testing and evaluation to assess their effectiveness, ecological impact, and safety before large-scale implementation.

Field trials represent the gold standard in evaluating mosquito control strategies. However, they require substantial financial resources, logistical support, and expert coordination to ensure proper implementation [20,21]. To date, trial results of inundative mosquito control strategies have shown high variability, highlighting a major knowledge gap in the underlying causes for this heterogeneity [9–18]. While pilot studies in Kentucky, Panama, Brazil, China, Cuba, and the Cayman Islands have yielded promising results in decreasing mosquito populations [10,12–15,17], other trials in Florida, Mexico, Italy, and Malaysia have had more limited success [9,11,16,22]. The lack of standardized guidelines on the design and implementation of field trials for inundative mosquito control strategies has possibly contributed to these results. For instance, the release ratio of sterilized, *Wolbachia*-infected, or genetically modified male mosquitoes to wild-type males varied widely [9,10,14], including trials that released mosquitoes without assessing the ratio with the wild-type population [13,15]. As another example, the trial start and end dates were variable between studies, with some trials being conducted during the low mosquito season [22], others during the high season [16], and some spanning most of the year [13,15].

Without standardized trial designs, it is challenging to compare findings across different studies and settings. Mathematical models have been used to inform and guide the implementation of field trials [23], including those for Ebola [24] and dengue [25] vaccines, and SARS-CoV-2 antiviral therapies [26,27]. Thus, our study introduces IRIS (Inundative mosquito Release Intervention Simulator), an agent-based model for the “in-silico” simulation of trial designs for inundative mosquito releases. As a case study, we focus on GMM-based strategies, evaluating two alternative trial designs to control the *Aedes aegypti* population, a primary vector for dengue, Zika, and chikungunya [28] and target species for previous trials [11,12,15]. As a study location, we selected Miami-Dade County, Florida, as it has an abundant *Ae. aegypti* population, an archive of historical mosquito surveillance data, and a history of local transmission of arboviruses [29–31].

## Methods

### Data

We used mosquito surveillance data from January 1, 2019, to December 31, 2023, for Miami-Dade County, FL, which is collected and managed by the Miami-Dade County Mosquito Control Division. The surveillance system consists of about 300 BG-Sentinel-2 traps baited with dry ice that are deployed once per week for 24 hours. Collected mosquitoes are identified using taxonomic keys. Details on the data collection are reported in our previous publication [32]. Daily mean temperature data for Miami-Dade County, Florida, was retrieved from publicly available records from the Miami International Airport [33].

### Model Formulation

As an illustrative example of our modeling tool, in this study, we simulate GMM-based trials that aim at reducing the mosquito population. GMM population suppression strategies involve the release of homozygous adult male mosquitoes carrying a gene that impairs the fertility of female offspring [34]. Specifically, when a genetically modified male mosquito mates with a wild adult female mosquito, it will produce a heterozygous offspring carrying that gene. Heterozygous females are infertile, while heterozygous males can mate with wild females, further propagating the gene in the wild mosquito population [19,34–37].

IRIS is an agent-based model that produces stochastic simulations of *Ae. aegypti* population dynamics considering four stages of mosquito development: egg, larva, pupa, and adult (Fig. 1A). The model estimates the number of *Ae. aegypti* in each stage of their life cycle for each calendar date at a time step of 0.1 days. Each mosquito is represented by an agent with the following attributes: sex (male or female), genetic makeup (genetically modified homozygous, genetically modified heterozygous, or wild type), and current stage of development (egg, larva, pupa, or adult). We assumed a 1:1 female-to-male ratio for eggs. At each time step of the simulation, each agent’s transition into the next development stage or mortality and adult female oviposition are sampled from a Bernoulli distribution using the appropriate temperature-dependent rate [38–41]. Wild-type adult female mosquitoes lay eggs at a given temperature-dependent oviposition rate [39,41] with an average of 100 eggs laid per oviposition [42]. Newly laid eggs then become new agents in the model. Eggs’ mortality is both temperature- and density-dependent to account for the carrying capacity of the environment (see section Model Calibration).

**Figure 1.**
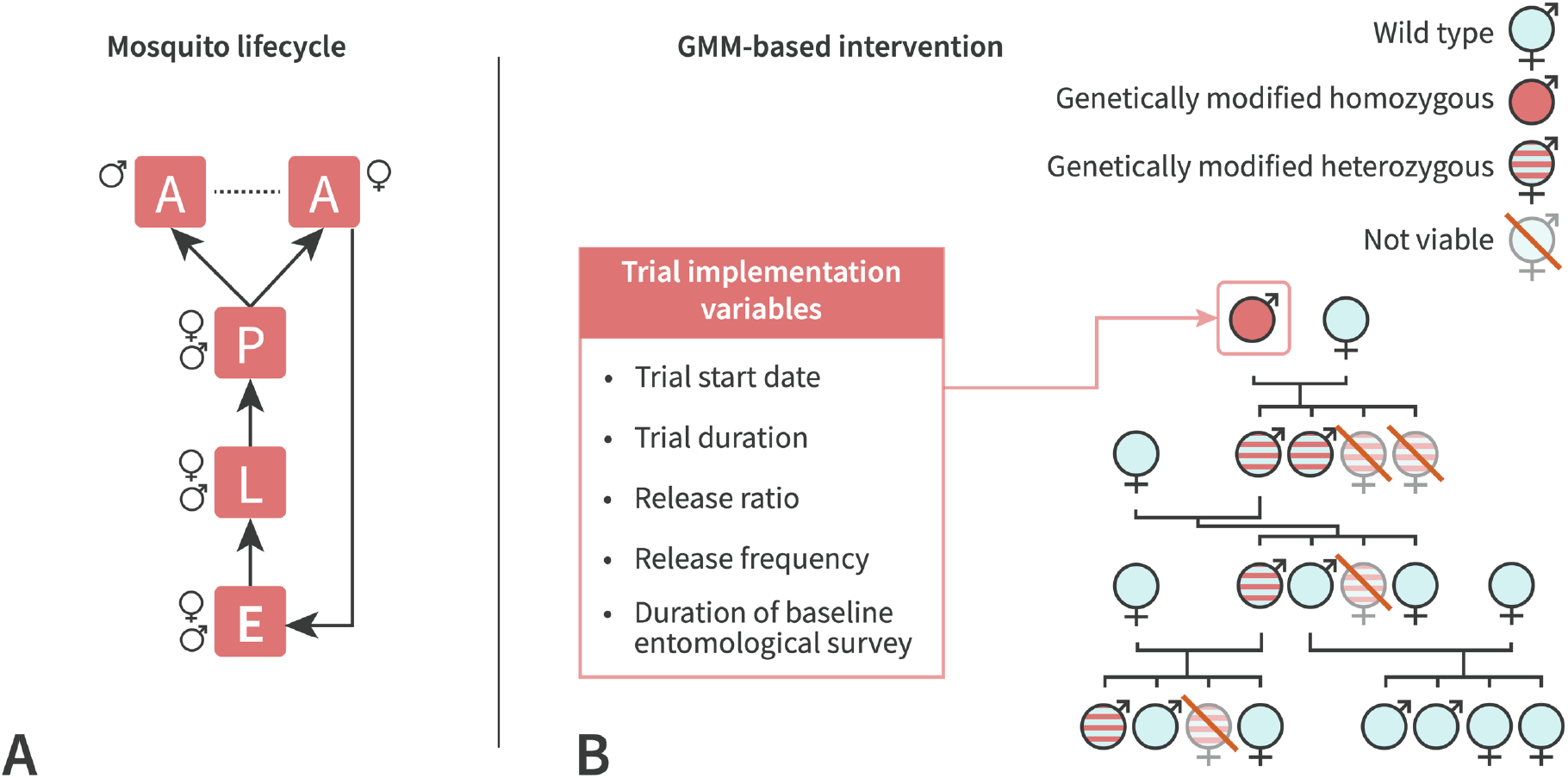
**A**. Conceptual representation of the modeled mosquito life cycle. **B**. Conceptual representation of the GMM mechanisms of action and list of trial design variables that are investigated in this study.

To simulate GMM-based trials, we periodically introduce a given number of new adult male agents that are genetically modified (homozygous) into the population. These mosquitoes continue through the life cycle in the same way as wild-type mosquitoes. When a wild-type female mates with a homozygous male, the offspring is heterozygous, and heterozygous females are infertile (i.e., their oviposition rate is set to 0). When a wild-type female mates with a heterozygous male, half of the offspring will be heterozygous for the transgene, and half will not carry the transgene. Moreover, we assume that female mosquitoes mate only once in their lifetime [43]. Finally, mosquitoes carrying the transgene have a different (generally lower) fitness than wild-type males, which reduces the likelihood of successful mating [44]. A conceptual representation of the process is shown in Fig. 1B.

### Model Calibration

Temperature-dependent development and mortality rates of mosquito lifecycle stages were taken from the literature [38–40]. The model also considers seasonal changes in the carrying capacity to account for variations throughout the year [41]. We used a Markov chain Monte Carlo (MCMC) procedure to obtain estimates of the posterior distributions of the carrying capacity for two periods of the year. MCMC explored the likelihood of observing the actual number of collected adult female *Ae. aegypti* specimens from Miami-Dade County, Florida, from a negative binomial distribution of the mean given by the model and the over-dispersion parameter, which is estimated as well [45]. Details are reported in the Supplementary Material.

### Simulation Analysis of Trial Implementation

The calibrated model simulates *Ae. aegypti* population dynamics from January 2019 through December 2023 and provides estimates of the daily number of *Ae. aegypti* specimens for each life stage of development, sex, and genetic makeup. This allows us to simulate different trial designs and trial implementation variables “in silico”. In this analysis, we simulated 2-arm trials where the control arm consists of the model realizations that are run without any GMM-based intervention, while the intervention arm consists of the model realizations simulating a GMM-based intervention.

We consider two trial designs:

- **Constant-release trial design.** The number of genetically modified (homozygous) male mosquitoes released per release in the intervention is set to obtain the desired ratio between the number of released male mosquitoes and the number of female mosquitoes per trap night collected during a baseline entomological survey. This baseline entomological survey is conducted before the start of the trial to assess the relative abundance of the mosquito population by monitoring traps for adult female mosquitoes (e.g., BG-Sentinel-2 traps). In this design, the number of released male mosquitoes per release is constant for the entire duration of the trial; thus, in populations following seasonal trends and where control is effective, the actual ratio of released male mosquitoes as compared to the actual female mosquito population changes over time.
- **Adaptive-release trial design**. The number of genetically modified (homozygous) male mosquitoes released per release in the intervention is dependent on the number of mosquitoes estimated in the control arm. The idea of the adaptive-release trial design is to rely on the control arm to obtain an estimate of the actual target female population relative abundance at any time point. Then, this estimate is used at the time of each release as the denominator to calculate the number of male mosquitoes to be released to obtain a constant release ratio as compared to the actual female mosquito population at any given point in time. Specifically, the number of male mosquitoes released at time *t, G*(*t*) is given by: 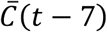, where *r* is the target release ratio, *s* is the trap collection ratio estimated in the literature (which corresponds to the ratio between the collected adult female mosquitoes and the total number of adult female mosquitoes), and 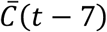 is the mean estimated number of mosquitoes in the control arm over the past week. The one-week lag was incorporated to account for the time required for trap collection, mosquito identification, data entry, and release planning.

In this study, the effectiveness of a simulated trial (trial outcome) is defined as the reduction in the *Ae. aegypti* female population within the monitored area during the trial period as:

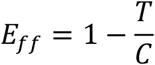

where *T* is the sum of estimated adult female mosquitoes in the treatment arm from 2 months after the start of the trial until the end of the trial, and *C* is the same quantity but for the control arm.

Results for each simulated trial are based on 100 stochastic model realizations obtained by uniformly sampling 100 indexes of MCMC corresponding to the joint posterior distributions of the carrying capacity for the two periods. Then, for each sampled index, we run the control and the treatment arm for both trial designs.

Standardized guidelines for trial implementation of GMM mosquito control strategies encompass a wide range of aspects and cover different steps of the implementation of the trial, from laboratory protocols to field practices. Among them, we investigate four variables: i) release ratio, ii) start date of the trial, iii) the duration of the baseline entomological survey (which is relevant only for the constant-release trial design), iv) release frequency, and v) trial duration. For the main analysis, we simulated 6-month trials with a release frequency of once per week, a 2-month baseline entomological survey, and that genetically modified mosquitoes have the same fitness as wildtype ones. We investigated release ratios from 0.1 to 9.0 by 0.1 steps, and we consider starting dates of the trial on the first day of the month for every month of the year in 2022. To estimate release ratios, we considered a 24.6% trap collection ratio within the trap catchment area [46]. Thus, an n:1 release ratio is relative to the collected female mosquitoes, which represents an underestimate of all female mosquitoes within the area.

We conducted several sensitivity analyses varying the release frequency (every 3, 7, and 14 days), trial duration (90, 180, and 365 days), duration of the baseline entomological survey (2, 6, and 12 months), and the fitness of the genetically-modified mosquitoes (set at 50% and 75% relative to the fitness of wild-type male mosquitoes). Moreover, we conducted a sensitivity analysis considering two alternative values for the collection ratio, 12.3% and 49.2%, encompassing the uncertainty reported in the reference study [46]. To assess possible differences between years, we conducted a sensitivity analysis where the trials were carried out in 2020, 2021, and 2022. Finally, to assess the impact of temperature and carrying capacity on the effectiveness of the GMM-based trial, we performed an analysis where the temperature was set to a constant 25°C and the carrying capacity was kept constant throughout the year, while all other parameters remained the same as in the main analysis.

## Code Availability

The codebase for IRIS is openly available on GitHub at https://github.com/jagadeesh-chitturi/iris-model.

## Results

The weekly number of adult female *Ae. aegypti* per trap night estimated by IRIS compares well with the observed data for Miami-Dade County over the entire study period (Fig. 2A). The data show a year-round presence of *Ae. aegypti* with a clear seasonal trend in its abundance, with peak activity taking place in July (about 10 collected adult female *Ae. aegypti* per trap night) and lower activity during the winter months, with about 2 collected specimens per trap night.

**Figure 2.**
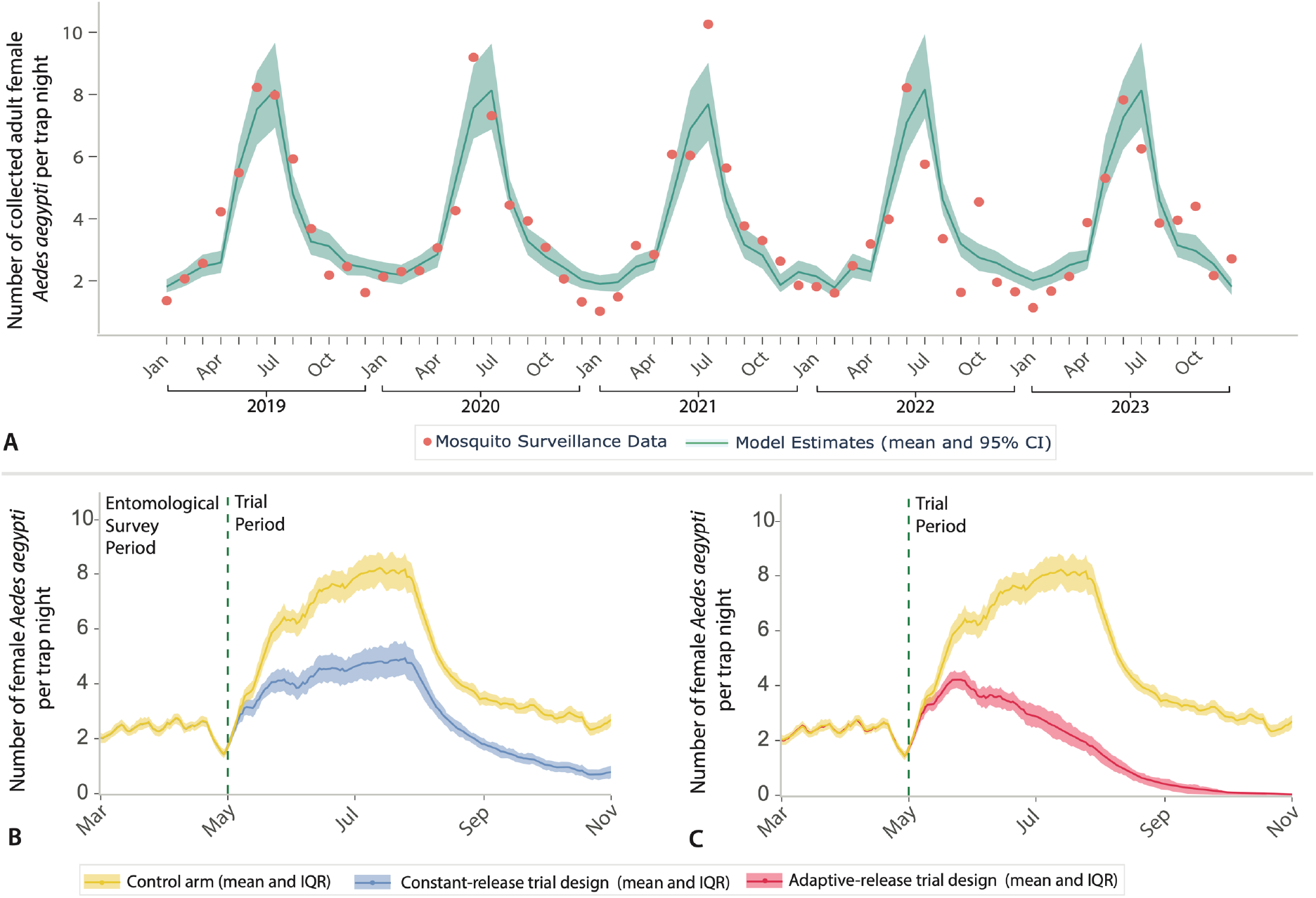
**A**. Number of collected adult female *Ae. aegypti* per trap night by month of the year as reported in the Miami-Dade mosquito surveillance system and as estimated by the model. **B**. Daily number of collected adult female *Ae. aegypti* per trap night estimated by the model in the control arm and in the treatment arm of a constant-release trial. Trial implementation details are: start date May 1, 6-month trial duration, weekly releases, 4:1 release ratio, and 2 months baseline entomological survey. **C**. As B, but for an adaptive-release trial design. Trial implementation details are the same as in panel B, except for the baseline entomological survey, which is not needed for this trial design.

Figure 2B shows an illustrative example of a trial simulated by IRIS: a constant-release GMM trial of a 6-month duration, weekly releases, a 4:1 release ratio, starting on March 1, and with a 2-month baseline entomological survey. Figure 2C shows the adaptive-release GMM trial based on the same implementation details as the constant-release trial design. The constant-release and adaptive-release trial designs follow a similar trajectory to the control arm for the first few weeks of the trial before diverging. Overall, we estimated an effectiveness of 47.3% (IQR: 37.5%-58.4%) and 81.3% (IQR: 71.3%-92.-0%) for the constant-release ratio and adaptive-release ratio, respectively. These results are not indicative of which trial design has a better suppression effectiveness; they merely represent the result of the considered implementation details and should not be interpreted as a comparative analysis of the two designs to identify an optimal one. For example, this simulated implementation of the adaptive-release trial requires a larger number of released mosquitoes over the entire trial period, which yields better effectiveness.

We used IRIS to explore different trial implementation details and their impact on the estimated effectiveness. First, we fixed weekly release frequency for a 6-month trial duration. For the constant-release design, we also fixed the duration of the entomological survey at 2 months. Regardless of the trial design, we found that the effectiveness is highly variable (Fig. 3A and B). For any trial start date, the effectiveness is highly dependent on the release ratio, which can lead to the outcome spanning the entire 0 to ∼100% range. If the release ratio is 12:1 or higher, the effectiveness is over 90% regardless of the trial start date. However, for lower values of the release ratio, the effectiveness is highly dependent on the trial start date (Fig. 3C). The estimated dependencies of trial effectiveness on the mosquito release ratio and trial start date may help explain the variability observed in field trials, which were based on different designs and implementations.

**Figure 3.**
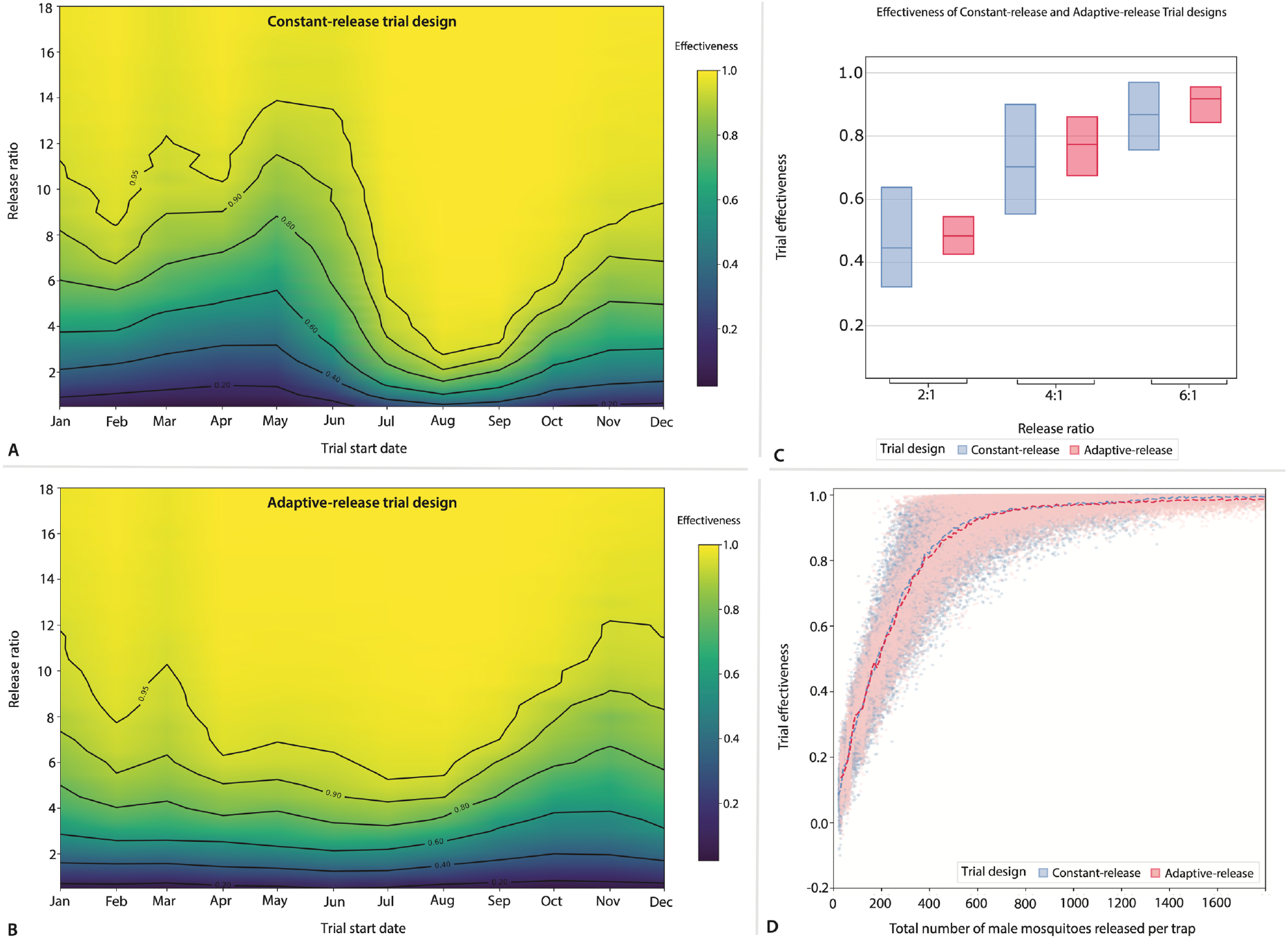
**A**. Estimated effectiveness of a constant-release trial as a function of the start date of the trial and release ratio. The other trial variables are set as follows: 6-month trial duration, weekly release, and 2-month baseline entomological survey. **B**. As A, but for an adaptive-release trial. **C**. Estimated median and IQR of the effectiveness of a constant-release and adaptive-release trial designs given a release ratio, for any possible trial start date. The other trial variables are set as in panel A. **D**. Estimated trial effectiveness as a function of the total number of genetically modified adult male mosquitoes released per trap over the entire duration of the trial. Each dot represents a single simulation; dashed lines represent the mean. The panel shows the results for 6-month trials with and weekly releases, any starting date, and any release ratio from 1:0.1 to 9:1. For the constant-release trial design, the duration of the baseline entomological survey is set to 2 months.

A more stable indicator we found to determine the effectiveness is the overall number of released mosquitoes during the entire duration of the trial (Fig. 3D). This holds true for any trial start date, release ratio, and trial design. In other words, a trial design with a given release ratio and start date requires a given number of mosquitoes to be released over the entire duration of the trial; this number essentially determines the final effectiveness of the trial.

Our results show that more frequent releases, longer trial durations, and greater fitness of transgenic mosquitoes increase the effectiveness of the trial and decrease the variability in the estimated effectiveness (Fig. 4A-C). For the constant-release design, a longer baseline entomological survey decreases variability (Fig. 4D).

**Figure 4.**
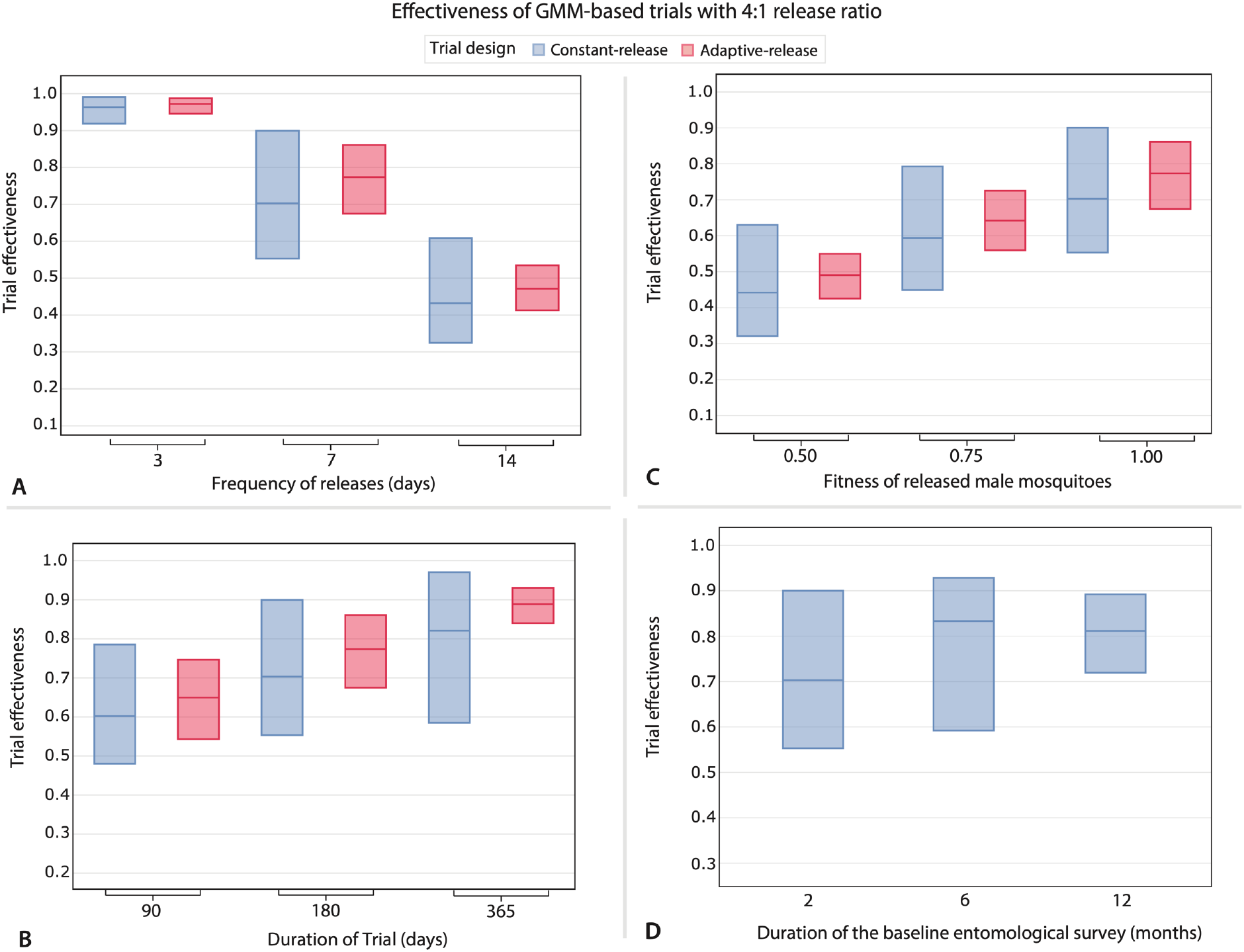
**A**. Estimated median and IQR of the effectiveness of a constant-release and adaptive-release trial designs given the release frequency, for any possible trial start date. The release ratio is set to 4 and the trial duration to 6 months; for the constant-release design, the duration of the baseline entomological survey is set to 2 months. **B**. Estimated median and IQR of the effectiveness of a constant-release and adaptive-release trial designs given the trial duration, for any possible trial start date. The release ratio is set to 4 and the release frequency to once per week; for the constant-release design, the duration of the baseline entomological survey is set to 2 months. **C**. Estimated median and IQR of the effectiveness of a constant-release and adaptive-release trial designs given the fitness of genetically modified male mosquitoes, for any possible trial start date. The release ratio is set to 4, the trial duration is set to 6 months, and the release frequency to once per week; for the constant-release design, the duration of the baseline entomological survey is set to 2 months. **D**. Estimated median and IQR of the effectiveness of a constant-release trial design given the duration of the baseline entomological survey, for any possible trial start date. The release ratio is set to 4, the trial duration is set to 6 months, and the release frequency to once per week.

We found that all our findings remain unchanged if trials are simulated using data from different years (see Supplementary Material). However, in a hypothetical scenario where the temperature and carrying capacity do not change over the course of the year, we estimated that a much larger release ratio is needed to obtain the same effectiveness estimated for the simulated trials using data for Miami-Dade County, FL (see Supplementary Material). Given that no changes take place over time and thus the mosquito population is constant year-round, the variability in the estimated effectiveness drastically decreases, suggesting that in areas where the mosquito population is more constant throughout the year, trial results may be more stable. Finally, we estimated a higher effectiveness of the trials for higher collection ratios of the traps, but our main finding about the lower variability in the effectiveness estimated using the adaptive-release design remains unchanged (see Supplementary Material).

## Discussion

In this study, we propose IRIS, an agent-based model designed to be a decision-support tool for public health authorities and researchers to in-silico test different trial designs and implementation details. This model builds on the wide body of existing literature on mathematical modeling for the evaluation of mosquito control strategies based on inundative mosquito releases [19,47–62], adapting it for trial simulation.

This analysis focuses on a specific study case: GMM-based trials for *Ae. aegypti* suppression in Miami-Dade County, Florida. Our results show that the effectiveness of GMM trials is highly sensitive to implementation choices, particularly the release ratio of modified males and the trial start date. This finding provides a possible explanation for the high variability observed in field trials across different settings [9–18]. Other trial implementation details, such as differences in release frequency, trial duration, and length of baseline entomological survey (for the constant-release trial design), may have further contributed to the observed variability in field estimates. On the other hand, our analysis suggests that the total number of modified mosquitoes released over the entire trial period may be a more stable indicator of trial effectiveness. This metric appears to transcend specific trial designs and implementation choices. Future research that considers a wider range of designs would be needed to further support/challenge this preliminary finding.

Our study is not designed to provide specific recommendations about field trial designs and their implementation details. Thus, we are not endorsing the implementation of a specific design. Instead, our work is intended to showcase a modeling approach for the in-silico simulation of trial designs and provide general insights to better interpret results from previous field studies. Our primary contribution is to provide an inexpensive and flexible modeling tool to test trial designs before committing the significant monetary and logistic resources needed for field implementation. This is especially relevant for proof-of-concept trials, which are the crucial first step in a pipeline toward full-scale public health adoption. The outcomes of such initial trials are critical, as they determine whether stakeholders may decide to commit resources to subsequent, larger-scale trials focused on optimizing long-term effectiveness (e.g., year-round release schedules). This optimization phase is, in turn, the prerequisite for the eventual, sustainable implementation of these strategies in real-world public health programs. IRIS directly supports the first step by informing the design of field trials with higher likelihood of success. Furthermore, IRIS codebase can be readily adapted to support both the design of larger-scale trials and the optimization of a wide range of other mosquito control interventions.

Our study is subject to the following limitations. First, our model simplifies certain biological, ecological, and environmental aspects. For example, it assumes a 1:1 sex ratio, female mosquitoes mating only once in their lifetime, uniform mixing between male and female mosquitoes, and relies on specific temperature-dependent parameters derived from the literature on *Ae. aegypti* rather than on the specifics of the population in the study site. While these assumptions are standard in many mosquito population dynamic models, they may not fully capture the nuances of real-world dynamics. Second, our analysis did not consider several logistical constraints and practical considerations associated with the implementation of field trials (e.g., implementation costs, required manpower, number of deployed traps per arm, number of clusters per arm, and the constant availability of a sufficient number of genetically modified male mosquitoes to release). However, although these constraints and considerations are crucial for the successful implementation of a trial, these should be considered by stakeholders to determine which designs and implementation details represent feasible options for them. Then, our tool can be used to compare these options to inform their decision among them. Third, our model was validated against historical mosquito surveillance data from Miami-Dade County, Florida, providing a good fit to observed population trends. However, it has not yet been validated against data from actual GMM field trials. Such validation would be crucial to confirm the model’s predictive power regarding the impact of GMM releases. Fourth, the current model does not account for spatial heterogeneity in breeding site availability, microclimatic conditions, etc., which may affect the outcome of a trial. Incorporating spatial factors into future model iterations would enhance the model’s capability to test more nuanced factors affecting the outcome of a trial. For example, the model currently considers a fitness parameter that represents the (potentially) reduced ability of genetically modified male mosquitoes to successfully mate. A spatial model would allow for the evaluation of different mechanisms driving such a reduction (e.g., shortened flight range, lower energy level) as well as boundary and spillover effects that are not considered here. Other differences between genetically-modified and wild-type male mosquitoes (e.g., lifespan, which is currently embedded into the fitness parameter) could be explicitly modeled and investigated as well. Fifth, our analysis focuses solely on GMM-based interventions and does not consider the potential for integrating other mosquito control methods. In practice, GMM releases might be combined with other interventions, such as the application of *Bacillus thuringiensis israelensis* (*Bti*) or other larvicides before the start of a trial.

In conclusion, this study presents a modeling tool designed for the “in-silico” simulation of field trials for mosquito control strategies based on inundative mosquito releases. Grounded in empirical mosquito surveillance data and incorporating key environmental factors, such as temperature-dependent development and mortality rates and temporal variations in carrying capacity, IRIS offers a platform for exploring different trial designs. While our analysis focused on *Ae. aegypti* in Miami-Dade County, our modeling approach allows for its adaptation to other mosquito species and geographic locations, making it an asset in the broader field of vector control. By enabling the identification of trial implementations with the highest likelihood of success, our model can contribute to the development of standardized guidelines for field trial design.

