## Supplementary Material for "IRIS, a tool for the in-silico evaluation of mosquito control trial designs based on inundative releases"

Jagadeesh Chitturi[1], Paulo C. Ventura[1], Allisandra G. Kummer[1], Chalmers Vasquez[2], Ethan SeRine[1], Megan D. Hill[1], Mattia Manica[3], Piero Poletti[3], John C. Beier[4], Keisuke Ejima[5], Michael Johansson[6], Stefano Merler[3], Hongjie Yu[7], John-Paul Mutebi[2], Maria Litvinova[8], André B.B. Wilke[8], Marco Ajelli[1]

[1]Laboratory for Computational Epidemiology and Public Health, Department of Epidemiology and Biostatistics, Indiana University School of Public Health, Bloomington, IN, USA

[2]Miami-Dade County Mosquito Control Division, Miami, FL, USA

[3]Center for Health Emergencies, Bruno Kessler Foundation, Trento, Italy

[4]Department of Public Health Sciences, Miller School of Medicine, University of Miami, Miami, FL, USA

[5]Lee Kong Chian School of Medicine, Nanyang Technological University, Singapore, Singapore

[6]Network Science Institute, Northeastern University, Boston, MA, USA

[7]School of Public Health, Key Laboratory of Public Health Safety, Ministry of Education, Fudan University, Shanghai, China

[8]Department of Epidemiology and Biostatistics, Indiana University School of Public Health, Bloomington, IN, USA

#### Table of Contents

*Methodology* ... *3*

The mosquito lifecycle and mating dynamics ... 3

Simulated release of genetically modified male mosquitoes ... 4

Baseline entomological survey ... 4

Model calibration ... 5

*Outcome Variability* ... *7*

*Sensitivity Analyses* ... *10*

Frequency of releases ... 10

Trial duration ... 11

Duration of the baseline entomological survey ... 12

Year of the intervention ... 13

Trial effectiveness under constant environmental conditions ... 14

Collection ratio ... 15

### Methodology

#### The mosquito lifecycle and mating dynamics

We developed an agent-based simulation to model the dynamics of mosquito populations. In the model, each agent represents an individual mosquito, which can be classified within stages of the lifecycle of a mosquito, including the aquatic stages – eggs, larvae and pupa – and the adult stage. Both male and female mosquitoes are included in the simulation. Male mosquitoes are further classified according to their genetic makeup, fitting into one of the categories: (i) genetically modified homozygous, which carry two copies of the modified gene; (ii) genetically modified heterozygous, which carry one copy of the dominant modified gene; (iii) wild-type, which do not carry any copy of the dominant modified gene. For simplicity, female mosquitoes in our simulation are always wild-type with respect to the genetic makeup, since female mosquitoes that carry any dominant modified gene copy are infertile. As female *Ae. aegypti* mate only once in their lifetime and store sperm in their spermathecae, we considered this mechanisms by adding a tag to simulated females that differentiate females that have already mated with those that did not.

The mosquito lifecycle is simulated as a progression between the lifecycle stages, including egg ($E$), larva ($L$), pupa ($P$) and adult mosquito ($A$), with stage-specific development and mortality rates. At each time step $\Delta t$ of the simulation ($\Delta t = 0.1$ days), we simulate the following processes:

An egg may develop into a pupa by sampling from a Bernoulli distribution with probability $d_e(T_t) \cdot \Delta t$, where $T_t$ is the temperature at time $t$ ($t$ represents a calendar day) and $d_e(T)$ is the temperature-dependent development rate for eggs at temperature $T$. An egg may also die and be removed from the simulation by sampling from a Bernoulli distribution with probability $m_e(T_t) \cdot \Delta t$, where $m_e(T)$ is the temperature-dependent mortality rate for eggs.

Similarly, a larva may develop into pupa with probability $d_l(T_t) \cdot \Delta t$ or die with probability $m_l(T_t) \cdot \Delta t$, where $d_l(T)$ and $m_l(T)$ are the temperature-dependent development and mortality rates for larvae, respectively. A pupa may develop into an adult mosquito with probability $d_p(T_t) \cdot \Delta t$ or die with probability $m_p(T_t) \cdot \Delta t$, where $d_p(T)$ and $m_p(T)$ are the temperature-dependent development and mortality rates for pupae.

An adult mosquito may die and be removed from the simulation with probability $m_a(T_t) \cdot T_t$, where $m_a(T)$ is the temperature-dependent mortality rate for adults. For adult mosquitoes, we also simulate mating and oviposition mechanics:

- The mating process is simulated as follows. At time step $\Delta t$, an adult female that has not already mated may mate based on the result of a random sample from a Bernoulli distribution with probability $d_a(T_t) \cdot \Delta t$, where $d_a(T)$ is the oviposition rate at temperature $T$. If the sampled value is 1, then her mate is randomly selected among all male mosquitoes. Two options are possible: 1) the selected male is wildtype, thus the probability of mating is 1; or 2) the selected male is genetically modified (either homozygous or heterozygous), thus we further sample from a Bernoulli distribution with probability $\varphi \leq 1$, representing the fitness parameter; if the sampled value is 1, then the mosquito mates, while if the sampled value is 0, the process repeats.

- The oviposition process is simulated as follows. Once a mate is found, the female mosquito may lay eggs (ovipose) with probability given by $1.0 - N_E(t)/K(t)$, where $N_E(t)$ is the number of eggs in the simulation and $K(t)$ is the carrying capacity at time $t$. If oviposition succeeds, up to $n_e$ new eggs are created in the simulation, distributed according to the following criterion following rules of Mendelian inheritance:
  - If the selected male adult is wild-type, $n_e/2$ wild-type male eggs and $n_e/2$ female eggs are created.
  - If the selected male adult is homozygous for the modified gene, only $n_e/2$ heterozygous male eggs are created.
  - If the selected male adult is heterozygous, then $n_e/4$ male heterozygous eggs, $n_e/4$ male wild-type eggs and $n_e/4$ female eggs are created.

#### Simulated release of genetically modified male mosquitoes

To simulate a field trial of the GMM method, the model includes release of homozygous adult male mosquitoes according to a predefined schedule, characterized by the start date, frequency, and duration. If a release is scheduled for time $t$, a number $G_t$ of new adult male homozygous mosquitoes are added to the simulation. The value of $G_t$ is defined depending on the selected release trial design:

- Constant-release trial design: A constant (i.e. time-independent) number of mosquitoes is released, $G_t = r \cdot s \cdot \overline{N_{A,f}}$, where $\overline{N_{A,f}}$ is the average number of collected mosquitoes on a baseline entomological survey (details below), $r$ is the ratio of released mosquitoes (e.g. 1:1, 2:1, 8:1) with respect to the surveyed population, and $s$ is the trap collection ratio.
- Adaptive-release trial design: The number of released mosquitoes is defined as $G_t = r \cdot s \cdot N_{A,f}(t - 7 \text{ days})$, where $N_{A,f}(t - 7 \text{ days})$ is the number of adult female mosquitoes in the simulation 7 days before the release time $t$. Variables $r$ and $s$ are respectively the release and collection ratios.

The first release occurs at the start date of the trial. Further releases are repeated every $n$ days (e.g., once every 3, 7 or 14 days), until the end date of the trial is reached.

#### Baseline entomological survey

For the constant-release trial design, we simulated a baseline entomological survey before the start of the trial. In the main analysis, we set the length of the baseline entomological survey to 2 months. For example, if the releases are scheduled to start on April 1, 2020, the entomological survey is conducted from February 1 to March 31, 2020.

During the entomological survey, the average number of *Ae. aegypti* female individuals is registered every week. Then we calculated $\overline{N_{A,f}}$ as the average of all weekly collections during the survey period, which is used as a baseline value for the constant release strategy as explained above.

#### Model calibration

The agent-based simulation model was calibrated to reproduce the monthly *Ae. aegypti* relative abundance data from Miami-Dade County, FL. We defined the likelihood of observing the number of collected adult female *Ae. aegypti* specimens given a model trajectory as:

$$\Lambda = \sum_{\{m \in M\}} \log\Big(\mathrm{NB}\big(F(m),\ M_f(m) \cdot O(m),\ \nu\big)\Big)$$

Where:

- $\mathrm{NB}(n, \mu, \nu)$ is the probability of observing $n$ from a negative binomial distribution of mean $\mu$, and over-dispersion $\nu$. The over-dispersion is explored as a free parameter in the calibration procedure.
- $F(m)$ is the empirical number of collected adult female *Ae. aegypti* specimens at month $m$.
- $M_f(m) = \sum_{t \in m} N_f(t)$ is the model-predicted abundance of *Ae. aegypti* specimens for month $m$, $N_f(t)$ is the number of female adults present in the

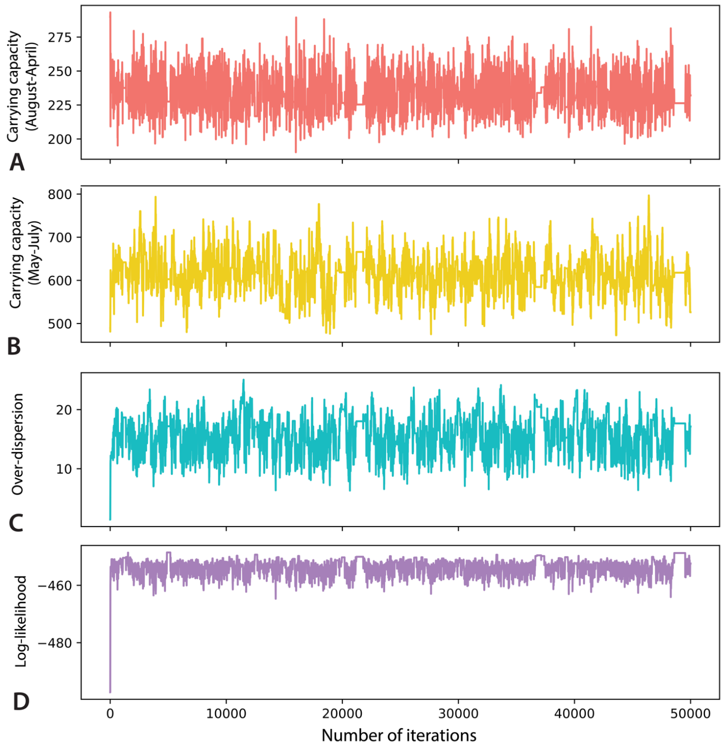


**Figure S1. A.** Traceplot of the estimated carrying capacity for August-April. **B.** As B, but for the carrying capacity for May-July. **C.** As A, but for the over-dispersion of the negative binomial distribution. **D.** As A, but for the log-likelihood.

simulation at the end of day $t$, and the sum in $t \in m$ runs through all days $t$ of month $m$.

- $O(m) = o(m)/(\max_{m \in M} o(m))$ is a normalized measure of the monthly trapping effort in Miami-Dade County, FL, where $o(m)$ is the total number of BG-Sentinel-2 trap deployments.

We explored the likelihood $\Lambda$ using Markov Chain Monte Carlo (MCMC) using the Metropolis-Hastings sampling algorithm with three free parameters: (i) the carrying capacity $K(t) = K_1$ for the period from August to April, (ii) the carrying capacity $K(t) = K_2$ for the period from May to July, and (iii) the overdispersion parameter $\nu$. The calibration chain consisted of 50,000 iterations (Fig. S1), from which the first 5,000 were considered as burn-in and discarded and the last 45,000 were used to estimate the posterior parameter distributions (Fig. S2).

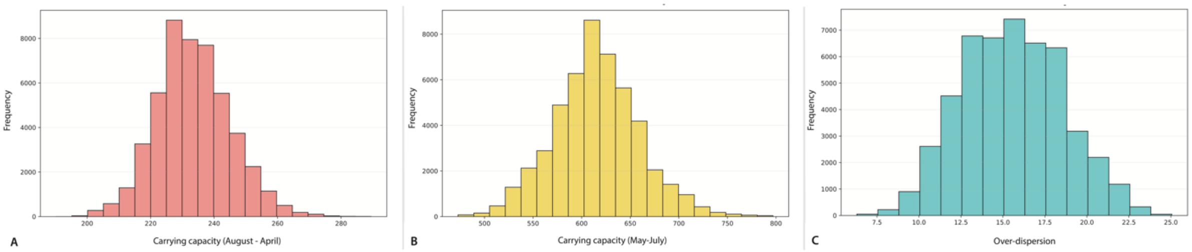


**Figure S2. A.** Estimated posterior distribution of the carrying capacity for August-April. **B.** As A, but for the carrying capacity for May-July. **C.** As A, but for the over-dispersion of the negative binomial distribution.

#### Outcome Variability

In Figs. 2B and 2C of the main text, we showed the mean and 95% confidence interval of the number of adult female mosquitoes by trap night during a trial. Here, we show randomly selected simulations, each one representing a single realization of a trial (Fig. S3 and S4).

We report the estimated 25% and 75% quantiles of the trial effectiveness (Fig. S5). The dependencies of trial effectiveness on the release ratio and trial start date follow the same trend as the estimated median shown in the main text.

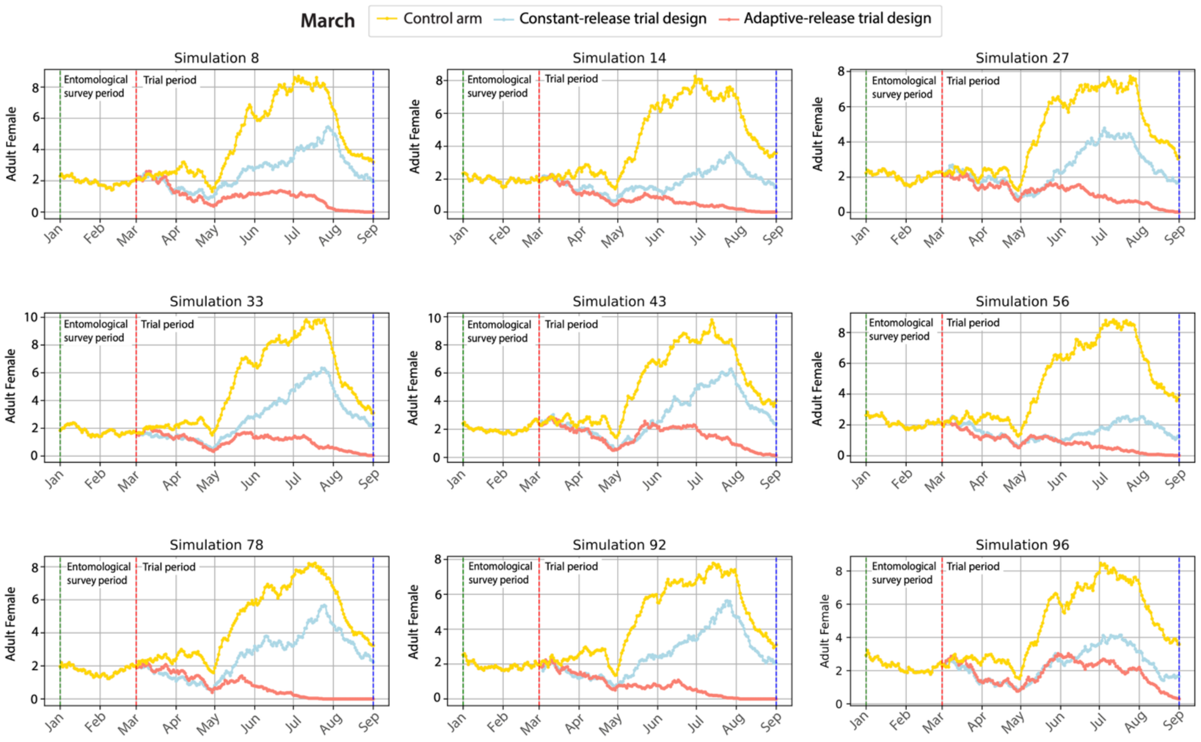


**Figure S3.** Daily number of collected adult female *Ae. aegypti* per trap night estimated by the model in the control arm and in the treatment arm of constant-release trial and an adaptive-release trial for nine randomly selected simulations. The trial duration is 6 months, releases are made weekly, the release ratio is 4:1, and the start date of the trial is March 1, 2022. For the constant-release trial, the duration of the baseline entomological survey is 2 months.

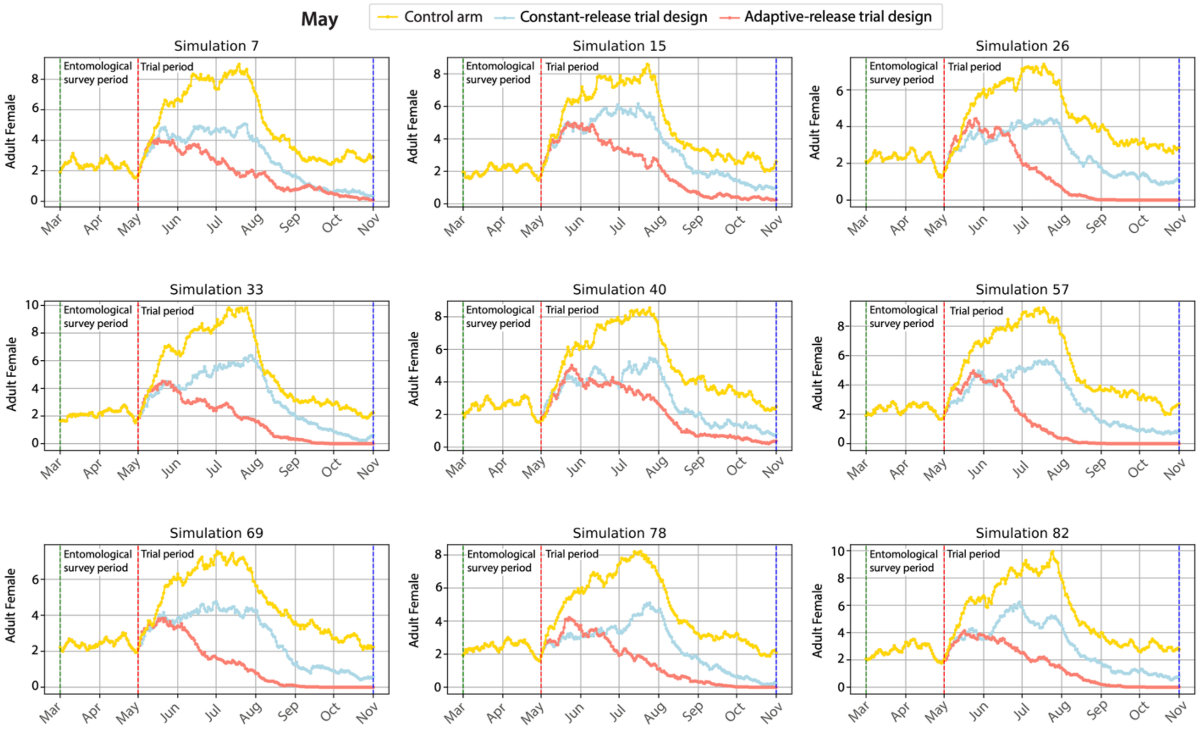


**Figure S4.** Daily number of collected adult female *Ae. aegypti* per trap night estimated by the model in the control arm and in the treatment arm of constant-release trial and an adaptive-release trial for nine randomly selected simulations. The trial duration is 6 months, releases are made weekly, the release ratio is 4:1, and the start date of the trial is May 1, 2022. For the constant-release trial, the duration of the baseline entomological survey is 2 months.

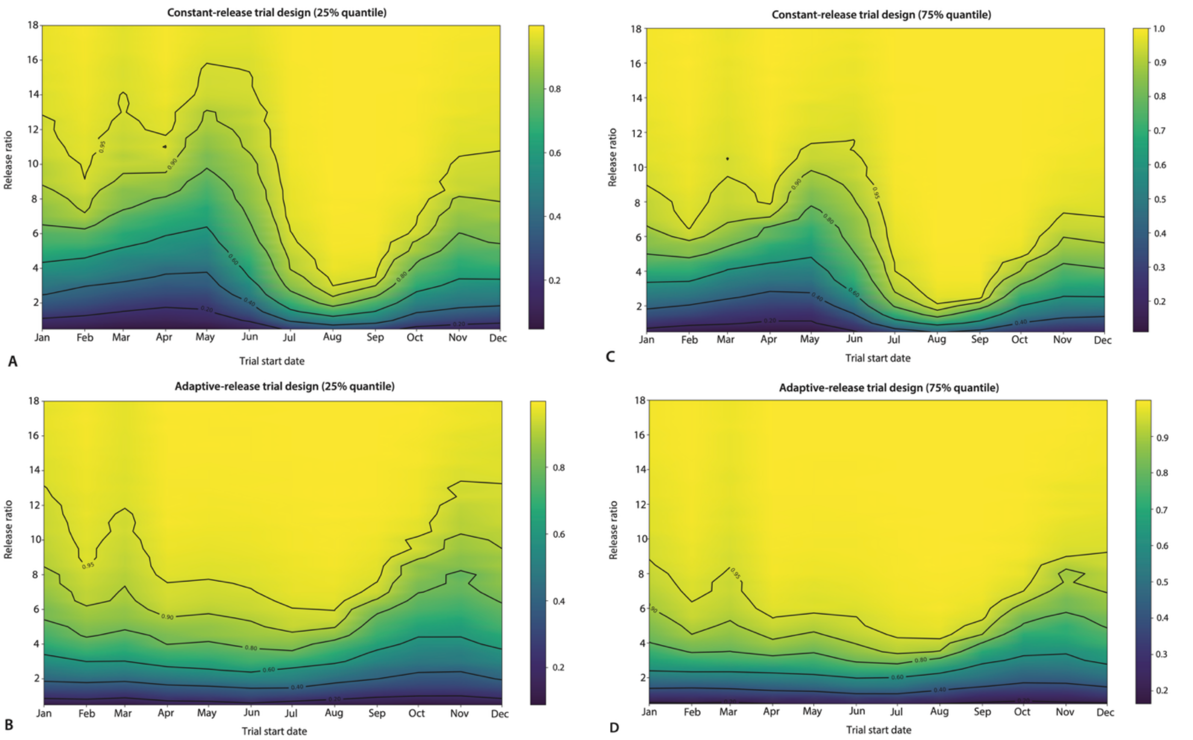


**Figure S5. A.** 25% quantile of the estimated effectiveness of a constant-release trial as a function of the start date of the trial and release ratio. The other trial variables are set as follows: 6-month trial duration, weekly release, and 2-month baseline entomological survey. **B.** As A, but for the adaptive-release trial design. **C.** As A, but for the 75% quantile for the constant-release trial design. **D.** As A, but for the 75% quantile for the adaptive-release trial design.

#### Sensitivity Analyses

##### Frequency of releases

In the main analysis, we used a release frequency of 7 days. Here, we conducted a sensitivity analysis using two alternative values: 3 and 14 days (Figs. S6). As we decrease the release frequency, we observe an increase in trial effectiveness and a decrease in outcome variability.

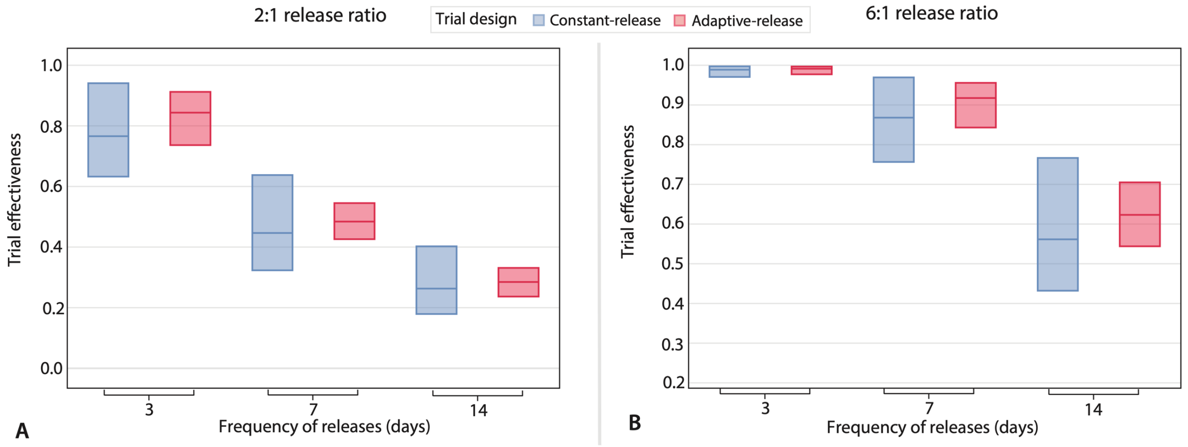


**Figure S6. A.** Estimated median and IQR of the effectiveness of a constant-release and adaptive-release trial designs given the release frequency, for any possible trial start date. The release ratio is set to 2:1 and the trial duration to 6 months; for the constant-release design, the duration of the baseline entomological survey is set to 2 months. **B.** As A, but for a 6:1 release ratio.

#### Trial duration

In the main analysis, we used a trial duration of 6 months. Here, we conducted a sensitivity analysis using two alternative values: 3 and 12 months (Figs. S7). As we increase the trial duration, we observe an increase in trial effectiveness and a decrease in outcome variability.

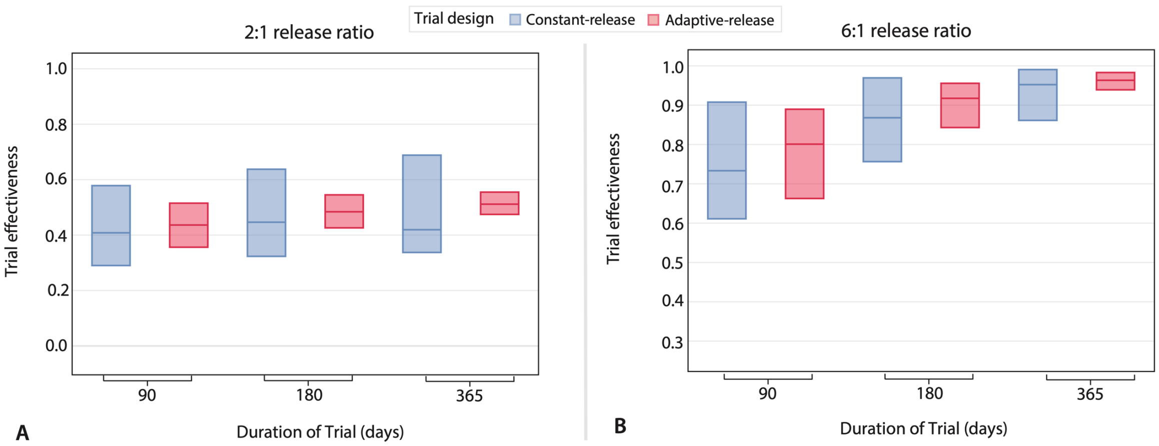


**Figure S7. A.** Estimated median and IQR of the effectiveness of a constant-release and adaptive-release trial designs given the trial duration, for any possible trial start date. The release ratio is set to 2:1 and the trial duration to 6 months; for the constant-release design, the duration of the baseline entomological survey is set to 2 months. **B.** As A, but for a 6:1 release ratio.

#### Duration of the baseline entomological survey

In the main analysis, we used a duration of the baseline entomological survey of 2 months. Here, we conducted a sensitivity analysis using two alternative values: 2 and 12 months (Figs. S8). While increasing the length of the baseline entomological survey has only a moderate effect on trial effectiveness, it reduces outcome variability.

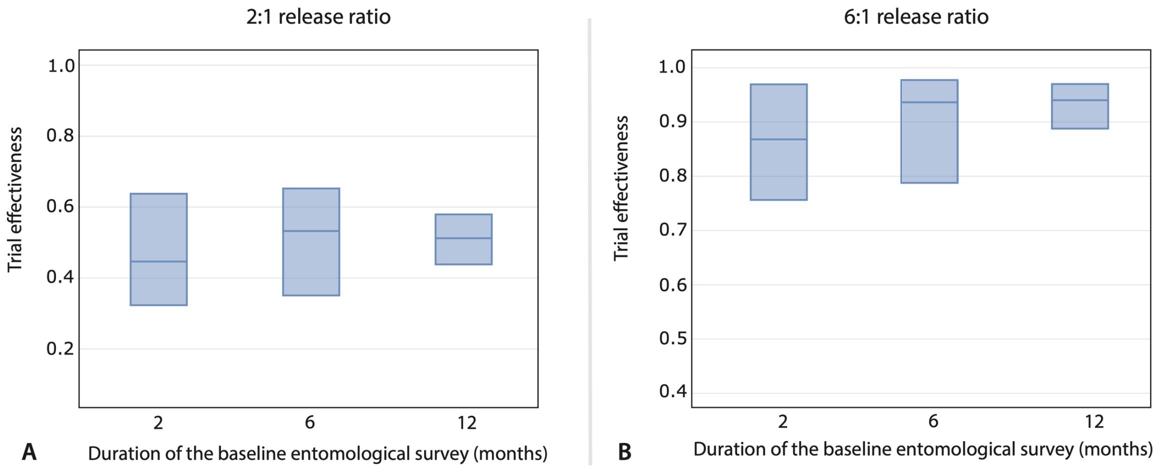


**Figure S8. A.** Estimated median and IQR of the effectiveness of a constant-release and adaptive-release trial designs given the duration of the baseline entomological survey, for any possible trial start date. The release ratio is set to 2:1 and the trial duration to 6 months; for the constant-release design, the duration of the baseline entomological survey is set to 2 months. **B.** As A, but for a 6:1 release ratio.

#### Year of the intervention

In the main analysis, we simulated trials in 2022, using Miami-Dade County temperature data for that year. Here we compare the main results with those obtained by simulating a trial in 2020 and 2021. Our results show that the estimated effectiveness of the trail is consistent between years (Fig. S9).

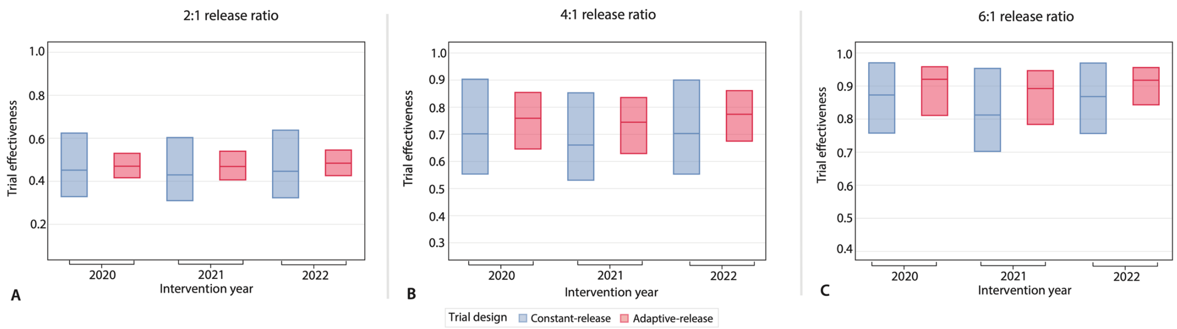


**Figure S9. A.** Estimated median and IQR of the effectiveness of a constant-release and adaptive-release trial designs given the intervention year, for any possible trial start date. The release ratio is set to 2:1 and the trial duration to 6 months; for the constant-release design, the duration of the baseline entomological survey is set to 2 months. **B.** As A, but for a 4:1 release ratio. **C.** As A, but for a 6:1 release ratio.

#### Trial effectiveness under constant environmental conditions

We conducted trial simulations under a hypothetical scenario with a constant temperature of 25ºC and a constant carrying capacity throughout the year. Results show remarkably less variability in the effectiveness for any release ratio when compared to the scenarios with time-dependent environmental conditions (Fig. S10 vs. Fig. 3 in the main text).

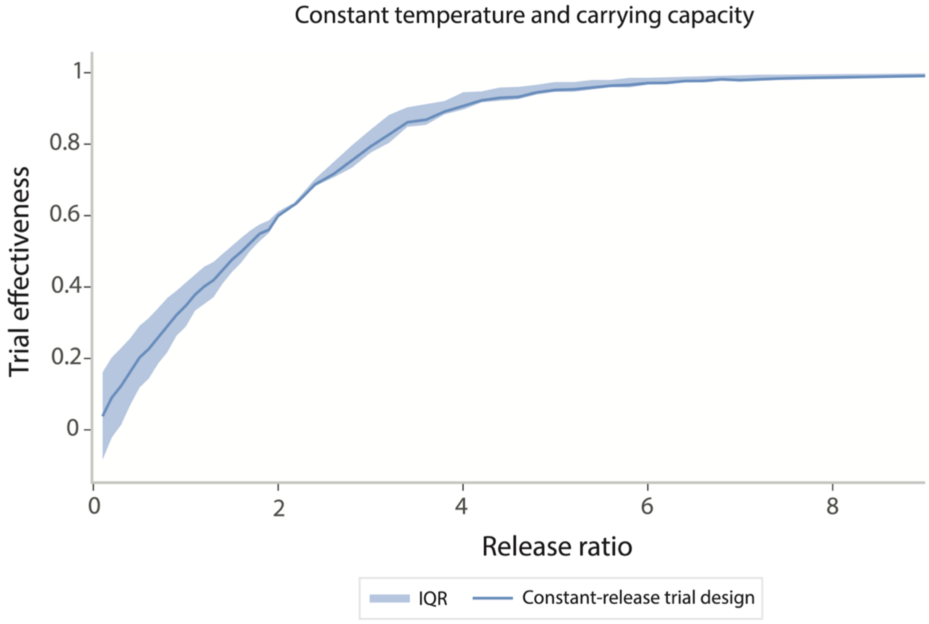


**Figure S10.** Estimated effectiveness of a 6-month constant-release trial design with weekly releases as a function of the release ratio under the assumption of a constant temperature and constant carrying capacity. The carrying capacity is set at the value estimated for the May-July period.

#### Collection ratio

Release ratios were set based on the number of collected mosquitoes, which only represent a fraction of the total mosquitoes within the trap catchment area. Here we considered two alternative values for the collection ratio: 2-times higher than the literature value used as baseline and 2-times lower, namely 49.2% and 12.3%. We found that a lower (higher) collection ratio value would mean that the actual population around the trap was underestimated (overestimated), implying in lower (higher) effectiveness with the same ratio. However, our finding about the variability of the estimate trial outcomes remains unchanged regardless of the collection ratio (Fig. S11).

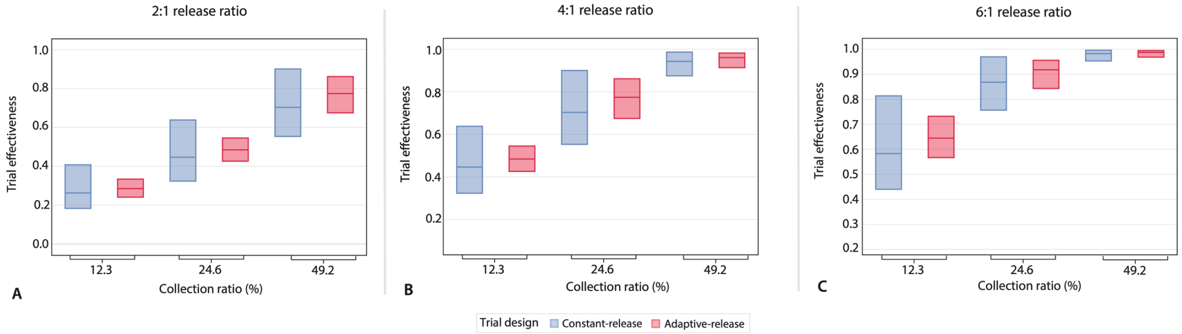


**Figure S11. A.** Estimated median and IQR of the effectiveness of a constant-release and adaptive-release trial designs given the collection ratio. The release ratio is set to 2:1 and the trial duration to 6 months; for the constant-release design, the duration of the baseline entomological survey is set to 2 months. **B.** As A, but for a release ratio set to 4:1. **C.** As A, but for a release ratio set to 6:1.
